# PPARγ/ETV2 Axis Regulates Endothelial-to-Mesenchymal Transition in Pulmonary Hypertension

**DOI:** 10.1101/2022.05.17.492293

**Authors:** Dong Hun Lee, Minseong Kim, Sarah S. Chang, Andrew J. Jang, Juyoung Kim, Jing Ma, Michael J. Passineau, Raymond L. Benza, Harry Karmouty-Quintana, Wilbur A. Lam, Roy L. Sutliff, C. Michael Hart, Changwon Park, Bum-Yong Kang

## Abstract

Endothelial-to-mesenchymal transition (EndoMT) plays an important role in pulmonary hypertension (PH). Also, the molecular mechanisms regulating EndoMT in PH remain to be defined. In this study, we first showed that reduced expression of the transcription factors ETV2 (ETS variant 2) and PPARγ (Peroxisome Proliferator-Activated Receptor gamma) along with reduced endothelial markers and increased EndoMT markers were consistently observed in lungs and pulmonary artery endothelial cells (PAECs) of idiopathic pulmonary arterial hypertension (IPAH) patients, in hypoxia-exposed mouse lungs, human PAECs, and in induced EndoMT cells. Base on this observation, we aimed to investigate the function of ETV2 and PPARγ in EndoMT. We have explored the function of ETV2 and PPARγ and its mechanism in PH using in *Etv2^+/-^* mice or PPARγ KO mice. *Etv2*^+/-^ mice spontaneously developed PH and right ventricular hypertrophy, associated with increased EndoMT markers and decreased EC markers. PPARγ transcriptionally activated the ETV2 promoter. Endothelial PPARγ expression in mice is positively correlated with ETV2 expression, but inversely with EndoMT markers.

Overexpression of ETV2 in hypoxia-exposed rat pulmonary artery led to vascular relaxation. We conclude that PPARγ-ETV2 signaling can function as a novel pathway in PH pathogenesis by attenuating EndoMT.

## INTRODUCTION

Pulmonary hypertension (PH), defined as an elevation of the mean pulmonary artery pressure >20 mmHg including a pulmonary vascular resistance (PVR) ≥3 wU (wood unit), causes high morbidity and mortality (1–3). PH is characterized by pulmonary endothelial dysfunction and abnormal proliferation of pulmonary vascular wall cells, vascular remodeling, and muscularization of small pulmonary vessels (4). These structural and functional alterations in the pulmonary vasculature increase pulmonary vascular resistance resulting in progressive right-sided heart failure and death.

Recent studies show that endothelial to mesenchymal transition (EndoMT) contributes to the pathogenesis of PH. EndoMT involves a phenotypic “switch” in which endothelial cells (ECs) lose endothelial and gain mesenchymal phenotypic features. This phenotypic and functional switch can ultimately lead to pulmonary vascular remodeling, which is characterized by extensive accumulation of cells expressing smooth muscle actin (αSMA) within the microvessels of the hypertensive lung and in fibroblasts (5, 6). Lineage tracking analysis in mice demonstrated that EndoMT contributes to a spectrum of structural changes including thickening of the adventitial, medial and intimal layers of the pulmonary artery (PA) wall, medial hypertrophy of muscular arteries, and muscularization of small pulmonary arterioles (7). Further, studies in a mouse PH model (8) and histological analysis of human patient samples (9) suggested EndoMT as a potential mechanism of distal pulmonary arteriole muscularization (9, 10). However, the molecular mechanisms controlling these events have yet to be fully elucidated and constitute the focus of the current study.

ETS Variant Transcription Factor 2 (ETV2), also known as ER71, belongs to the ETS transcription factor family classified by the presence of the conserved ETS DNA binding domains. ETV2 functions as a critical regulator of the vertebrate cardiovascular system (10). We have demonstrated that *Etv2* deficient mouse embryos exhibited complete absence of the embryonic vasculature and died in utero as early as embryonic day 10.5 (11). Similar results were reported from other studies using alternative strategies such as gene trap and knockin/knockout approaches (12, 13). Given the conserved functions from zebrafish (14) and *Xenopus er71* (15), it is clear that ETV2 is indispensable for cardiovascular development. We have also shown that endothelial *Etv2* is also required for new vessel formation in response to injury and that delivery of *Etv2* into ischemic hindlimbs promotes perfusion recovery with concomitant neovascularization (16). Further, studies including ours have revealed that ETV2 alone is sufficient to convert human dermal fibroblasts (HDFs) into functional ECs (17, 18). Taken together, these findings strongly suggest that ETV2 is a master regulator of EC fate.

PPARγ (Peroxisome Proliferator-Activated Receptor gamma) is a ligand-activated transcription factor of the nuclear hormone receptor superfamily (19). Its expression is reduced in the pulmonary vasculature of patients with severe PH (20). Furthermore, EC-targeted depletion of PPARγ caused spontaneous PH in mice (21). In contrast, activation of PPARγ with thiazolidinedione (TZD), PPARγ agonists attenuates PH or vascular remodeling in essentially every experimental model of PH in which they have been tested (22). These results suggest a pivotal role for PPARγ in the pathogenesis of PH. In this study, we investigated novel functions of the of PPARγ-ETV2 axis in regulating EndoMT in PH.

## MATERIALS AND METHODS

### Control and IPAH lung tissues and PAECs

We obtained peripheral lung tissues from control or idiopathic pulmonary arterial hypertension (IPAH) patient specimens collected by the Pulmonary Hypertension Breakthrough Initiative (PHBI). IPAH samples were derived from 2 male and 3 female patients, 24-56 years old whereas control specimens were derived from 2 male and 3 female subjects, 29-55 years old who were failed donors. Human PAECs isolated from the lungs of control or IPAH subjects as described (23) were generously provided by Dr. Harry Karmouty-Quintana (University of Texas Health Science Center at Houston, Houston, TX).

### *In vivo* mouse model of PH

Male C57BL/6J mice, aged 8-12 weeks, were treated three times with weekly injections of the VEGF receptor antagonist, Sugen 5416 (SU, 20 mg/kg, subcutaneous injection) and exposed to hypoxia (HYP/SU, 10% O_2_) or normoxia (NOR/SU, 21% O_2_) for 3 weeks as reported (24). To assess PH, right ventricular systolic pressure (RVSP) and right ventricular hypertrophy (RVH) were measured in NOR/SU and HYP/SU-treated mice and in male *Etv2*^+/-^ mice as reported (24). All animals were given unrestricted access to water and standard mouse chow. All animal studies were approved by the Institutional Animal Care and Use Committee of Emory University or the Atlanta Veterans Affairs Healthcare System.

### *In vitro* cell models

For the hypoxia experiment with ECs, HPAECs were exposed to NOR or HYP (1% O_2_) conditions for 72 hours as reported (25). To induce EndoMT, HAPECs (passage 2-6, ScienCell, Carlsbad, CA) were incubated with 0.1 ng/mL interleukin-1 beta (IL-1β), 10 ng/mL tumor necrosis factor alpha (TNFα), and 10 ng/mL transforming growth factor beta (TGFβ) (*i*-EndoMT) or DMSO (CON) for 72 hours. Levels of HPAEC mRNAs associated with EndoMT were determined using qRT-PCR.

### Luciferase-based promoter assay

HEK/293T cells (1.3×10^5^/well of a 24-well plate) were transfected with 2 μg PPARγ expression plasmid (pcDNA3.1-FLAG-PPARγ), 200 ng pGL3-ETV2 promoter (26), and 30 ng pRL-null by lipofectamine 2000 (Thermo Fisher Scientific, Waltham, MA). Forty-eight hours later, cells were harvested, and luciferase activity was measured using the Dual-Luciferase reporter assay system (Promega, Madison, WI) according to the manufacturer’s instructions. Firefly luciferase values were divided by Renilla luciferase values to normalize transfection efficiency. Rosiglitazone (10 μM) was added into the culture 12 hours prior to cell harvest.

### Scratch wound assay

Human pulmonary artery endothelial cells HPAEC (3 × 10^5^/well) derived from 3 separate individuals were cultured in 6 well plates with ECM media (ScienCell, Carlsbad, CA) then treated with TGF-β (10 ng/ml), TNF-α (10 ng/ml) and IL-1β (0.1 ng/ml) for 24 hours or 72 hours. Then, treated cells (2.8 × 10^4^/well) were cultured overnight to reach a confluent layer in a well separated by an insert (Culture–Insert 2 Well, ibidi GmbH, Germany). Subsequently, the insert was removed to generate the cell-free gap and add 1 ml of ECM media (ScienCell, Carlsbad, CA). Images were taken at 0 and 5 hours after incubation using a phase-contrast microscope. The area of the cell-free gap was calculated by Image J.

### Immunohistochemistry and Immunocytochemistry

For each lung, sections from 10 separate tissue blocks were stained with hematoxylin and eosin for immunobiological analysis of pulmonary tissue. Cells in a slide glass were fixed with 4% paraformaldehyde and washed with 1x PBS for 5 times. Subsequently, cells were incubated with anti-ETV2, anti-SLUG, anti-TWIST, anti-DESMIN, anti-SMA, or anti-Ve-CDH5 antibody, followed by anti-mouse Alexa 488 or anti-rabbit 567 antibody. The images were taken using Olympus IX51.

### Collagen gel cell contraction assays

Suspension of *i*-EndoMT cells (5×10^5^) was mixed with collagen solution in a 1:4 ratio of cell suspension and collagen mixture provided in the cell contraction assay (Cell Biolabs) according to the manufacturer’s instructions. The gels were imaged at 0h and 120 h later and analyzed by ImageJ software.

### ETV2 or PPARγ gain and loss of function

For ETV2 or PPARγ loss of function, HPAECs were transfected with scrambled or *ETV2* or *PPARγ* RNAi duplexes (10 and 20 nM, Integrated DNA Technologies, Coralville, IA) using Lipofectamine 3000 transfection reagent (Invitrogen) according to the manufacturer’s instructions. After transfection for 6 hours, the transfection media were replaced with EGM containing 5% FBS and incubated at room temperature for 72 hours. HPAEC lysates were then harvested and examined for *PPARγ*, *ETV2*, *SLUG*, *TWIST1*, *DES*, *FSP1*, *αSMA*, *PECAM1/CD31*, and *VE-Cad/CDH5* levels using qRT-PCR analysis. To overexpress ETV2, HPAECs were transfected with ETV2 plasmid constructs (1-2.5 μg, oxETV2) or empty vector. For overexpression of PPARγ, HPAECs were transfected with adenovirus containing a PPARγ plasmid (AdPPARγ, 25-50 multiplicity of infection, MOI) or control GFP plasmid as we previously reported (27). After transfection for 6 hours, media were replaced with fresh 5% FBS EGM, and HPAEC were then treated with normoxia (NOR, 21% O_2_) or hypoxia (HYP, 1% O_2_) for 72 hours. HPAEC lysates were then harvested and examined for *PPARγ*, *ETV2*, *SLUG*, *TWIST1*, *DES*, *FSP1*, *αSMA*, *PECAM1/CD31*, and *VE-Cad/CDH5* levels using qRT-PCR analysis.

### ETV2 overexpression in hypoxia-exposed rat pulmonary artery

Secondary branch pulmonary artery segments, 3 mm in length, were isolated from hypoxia/sugen-treated Sprague-Dawley rat lungs (age 8-10 weeks) for 3 weeks and reoxygenation for 2 weeks and exposed to control conditions or 1 × 10^6^ ad*ETV2* for 24 hours. Following incubation, arteries were mounted on stainless steel wires, placed in an organ chamber containing Krebs-Henseleit buffer and connected to a Harvard apparatus differential capacitor force transducer. For each artery, resting tension was set to 10 mN the lowest resting tension that maximum contraction to 50mM KCl was observed. Data are recorded using PowerLab digital acquisition and analyzed using Chart Software. Results are expressed as mean + SEM. Concentration-response curves were generated to the contractile agonist phenylephrine (PE, 0.1 nM to 10 μM) and ED_80_ determined. Studies examining endothelium-dependent relaxation were carried out by measuring responses to methacholine (MCh; 1 nM to 10 μM) and sodium nitroprusside (SNP; 0.1 nM to 1 μM) as previously described (28, 29).

### mRNA quantitative real-time polymerase chain reaction (qRT-PCR) analysis

To measure *PPARγ*, *ETV2*, *SLUG*, *TWIST1*, *DES*, *FSP1*, *αSMA*, *PECAM1/CD31*, and *VE-Cad/CDH5* levels, total RNAs in IPAH lungs, IPAH ECs, HPAECs, mouse lungs or *i*-EndoMT cells were isolated using the mirVana kit (Invitrogen). *PPARγ*, *ETV2*, *SLUG*, *TWIST1*, *DES*, *FSP1*, *αSMA*, *PECAM1/CD31*, and *VE-Cad/CDH5* mRNA levels in the same sample were determined and quantified using specific mRNA primers as previously described (8). *GAPDH* mRNA levels were used as a control.

### Western blot analysis

After treatment with rosiglitazone or vehicle, protein homogenates from mouse lungs or HPAECs were subjected to Western blot analysis as reported (8). Primary antibodies purchased from Santa Cruz Biotechnology (Santa Cruz, CA) included: SLUG mouse monoclonal antibody (1:500 dilution, Cat # SC-166476, 30 kDa), TWIST1 mouse monoclonal antibody (1:500 dilution, Cat # SC-81417, 28 kDa), DES mouse monoclonal antibody (1:500 dilution, Cat # SC-23879, 53 kDa), PECAM1 rabbit polyclonal antibody (1:500 dilution, Cat # SC-8306, 130 kDa) and VE-Cad rabbit polyclonal antibody (1:500 dilution, Cat # SC-28644, 130 kDa). Primary FSP1 mouse antibody (1:1000 dilution, Cat # 07-2274, 12 kDa) was purchased from EMD Millipore (Burlington, MA). Primary αSMA rabbit polyclonal antibody (1:500 dilution, Cat # RB-9010-PO, 42 kDa) was purchased from Thermo Scientific (Waltham, MA). GAPDH rabbit polyclonal antibody (1:10,000 dilution, Cat # G9545, 37 kDa) was purchased from Sigma-Aldrich (St. Louis, MO). Relative protein levels were visualized using Li-Cor proprietary software, quantified Image J software, and normalized to GAPDH levels within the same lane.

### Statistical Analysis

All data are presented as mean ± standard error of the mean (SE). Data were analyzed using analysis of variance (ANOVA). Post hoc analysis used the Student Neuman Keuls test to detect differences between specific groups. In studies comparing only two experimental groups, data were analyzed with Student’s t-test to determine the significance of treatment effects. The level of statistical significance was taken as p<0.05. Statistical analyses were carried out using GraphPad Prism, Version 8.0 software (LaJolla, CA).

## RESULTS

### ETV2 expression is downregulated in PH *in vivo* and *in vitro*

We first examined the expression of markers of EndoMT and ECs in lungs and pulmonary artery ECs (PAECs) isolated from patients with IPAH. As shown, the expression of EndoMT markers including *SLUG*, *TWIST1*, *DES*, *FSP1*, and *αSMA* was increased in lungs (**Figure 1A**) and PAECs (**Figure 1B**) isolated from patients with IPAH, whereas the expression of endothelial markers, *PECAM1/CD31* and *VE-Cad/CDH5* was decreased. A similar finding was observed in mice following the induction of PH with hypoxia and the VEGF receptor antagonist, Sugen 5416 (HYP/Su) treatment (**Figure 1C**). Hypoxia was also sufficient to increase EndoMT markers and decrease endothelial markers in human PAECs (HPAECs) in vitro (**Figure 1D**). Previous studies showed that lack of *Etv2* leads to a complete absence of ECs (11–13) and forced expression of *ETV2* converted non-ECs into ECs (17, 18) suggesting that ETV2 has a determinant role for regulating EC functions. Thus, we hypothesized that the expression of ETV2 is downregulated in PH lungs leading to the transition of EC to mesenchymal cells. Interestingly, we found that *ETV2* levels were significantly reduced in both lungs (**Figure 1E**) and PAECs (**Figure 1F**) isolated from IPAH patients, in lung tissues from HYP/Su mice (**Figure 1G**), and in hypoxia-exposed HPAECs (**Figure 1H**) suggesting a potential role of ETV2 in PH.

**Figure 1.**
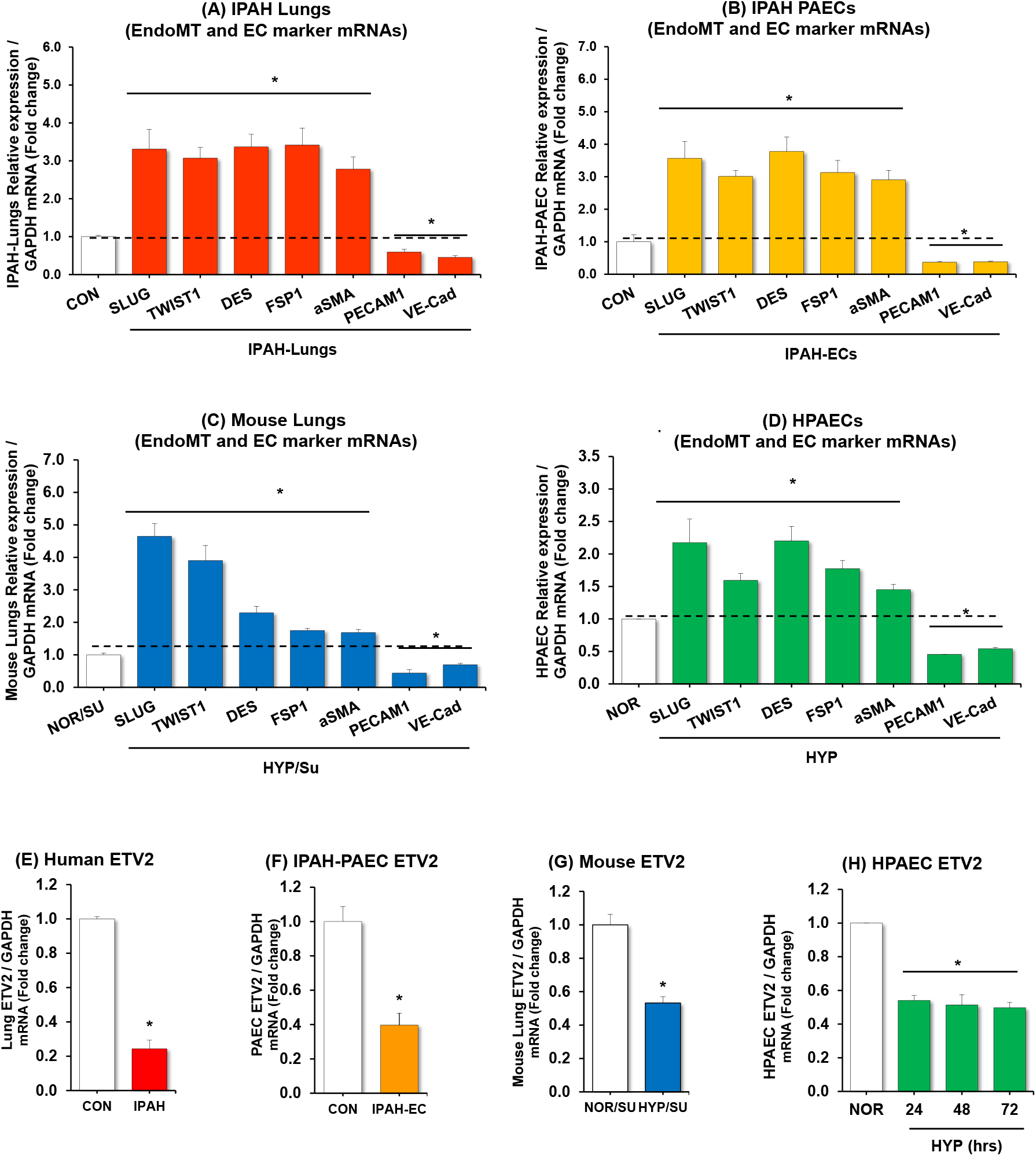
Expression of ETV2 decreases in PH patients and in mice upon hypoxia. **(A-B, and E-F)**, RNAs isolated from levels of IPAH lungs (A, E) or IPAH PAECs (B, F) were subjected to qRT-PCR. The results were expressed relative to *GAPDH* mRNA ± SE as fold-change vs. CON. *p<0.05 vs NOR, n=5. **(C and G)**, whole lung or pulmonary artery homogenates were collected from normoxic (21%) with sugen (NOR/SU, 20 mg/kg) or hypoxic (10% O_2_)/sugen (HYP/SU) mice for 3-weeks. Levels of lung EndoMT, EC marker, and *ETV2* were measured with qRT-PCR and expressed relative to lung *GAPDH* mRNA *p<0.05 vs NOR, n=6. **(D and H),** HPAECs (control) were exposed with normoxia (NOR, 21% O_2_) or hypoxia (HYP, 1% O_2_) for 72 hours. Mean levels of HPAEC EndoMT, EC marker, and ETV2 were measured with qRT-PCR. All bars represent the mean EndoMT or EC marker mRNA levels relative to *GAPDH* ± SE expressed as fold-change vs. NOR. *p<0.05 vs. NOR, n=6.

### Reductions in ETV2 stimulate EndoMT markers and PH *in vivo*

To determine the function of ETV2 in the pathogenesis of PH, we examined whether loss of ETV2 can lead to the development of PH. Since *Etv2^-/-^* mice are embryonic lethal (11), we used *Etv2^+/-^* mice which are fertile, normal in appearance and behavior, and display no overt overall or vascular phenotype (16). However, we found that *Etv2^+/-^* mice (aged 12-16 weeks) developed mild PH (**Figure 2A**) assessed by increased right ventricular systolic pressure (RVSP) and right ventricular hypertrophy (RVH) (**Figure 2B**). Importantly, enhanced positivity of αSMA, a indicative of vascular remodeling (**Figure 2C**) in *Etv2^+/-^* mice. Compared to littermate controls, the expression of EndoMT markers was increased and endothelial markers were reduced in the lungs of *Etv2^+/-^* mice (**Figure 2D and E**). Further, we performed siRNA-mediated *ETV2* depletion in HPAEC and found induction of expression of EndoMT markers and reduction of endothelial markers in the cells (**Figure 2F and G**). Lentiviral-mediated-*ETV2* overexpression in HPAECs led to downregulation of the EndoMT markers and was able to increase the expression of *PECAM1* and *VE-Cad/CDH5* in HPAECs under hypoxic conditions (**Figure 2H and I**). Collectively, these results suggest that endothelial ETV2 could have functional significance in PH.

**Figure 2.**
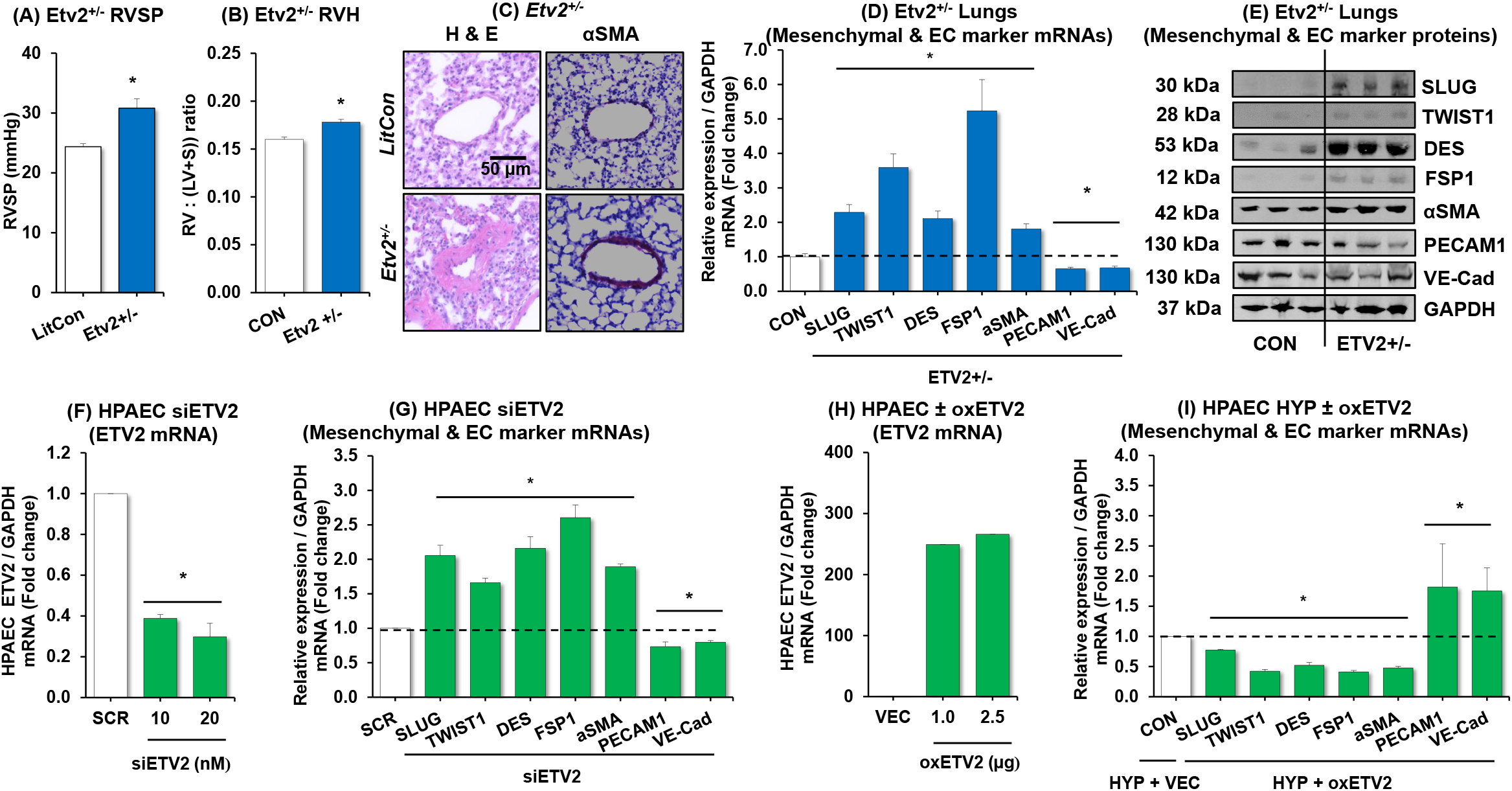
Reduction of ETV2 leads to the development of PH with increased expression of mesenchymal markers. **(A)** Right ventricular systolic pressure (RVSP) was recorded in anesthetized mice with a pressure transducer. Each bar represents the mean RVSP in mmHg ± SE. n=5-8. **(B)** The ratio of the weight of the right ventricle to the left ventricle + septum [RV: (LV + S)] is presented as an index of right ventricular hypertrophy (RVH). n=5-8. **(C)** Representative images of small mouse arterioles following H&E or αSMA staining. **(D-E)** Levels of lung EndoMT or EC markers were measured with qRT-PCR (C) or with Western blotting (D). The qRT-PCR results were expressed relative to *GAPDH* mRNA or protein ± SE as fold-change vs. CON. *p<0.05 vs NOR, n=5-7. **(F-G)** For ETV2 knockdown experiments, HPAECs were treated with scrambled (SCR) or *ETV2* (10-20 nM) siRNAs for 6 hours then incubated for an additional 72 hours. **(H-I)** For *ETV2* gain of function, HDF or HPAECs were treated with ETV2 plasmid constructs (oxETV2, 1-2.5 μg) or vector (VEC) constructs for 6 hours, then incubated for an additional 72 hours of normoxia (NOR) or hypoxia (HYP) exposure. qRT-PCR was performed for *ETV2*, EndoMT, or EC markers. Each bar represents mean ± SE *ETV2* or EndoMT or EC markers level relative to *GAPDH* expressed as fold-change vs cells treated with SCR. n=3-6, *p<0.05 vs VEC or NOR/VEC(-). +p<0.05 vs HYP/VEC(-).

### PPARγ functions as a direct upstream regulator of ETV2 expression

Studies have demonstrated that activation of the nuclear hormone receptor PPARγ attenuates PH whereas loss of PPARγ promotes PH (22). As illustrated in Figure E1 in the online data supplement, PPARγ levels were reduced in lungs and PAECs isolated from IPAH patients, in HYP/SU mouse lungs, and in hypoxia-exposed HPAECs. As shown in **Figure 1 and 2**, the expression of ETV2 is well correlated with pathogeneic features of PH, suggesting a potential role for the PPARγ and ETV2 axis in PH. Consistent with this postulate, our *in silico* analysis identified two putative PPREs (PPARγ Responsive Elements) within the *ETV2* promoter region (NM_014209, data not shown). To determine whether PPARγ can transcriptionally activate the expression of ETV2, we performed a luciferase-based ETV2 promoter assay. As shown in

**Figure 3**, overexpression of PPARγ in HPAECs using adenovirus-mediated *PPARγ* transduction (AdPPARγ, **Figure 3A**) caused a significant increase in *ETV2* expression (**Figure 3B**). In addition, AdPPARγ enhanced the activity of the *ETV2* promoter (**Figure 3C**). The observed promoter activity was further increased upon treatment with the PPARγ ligand, rosiglitazone (RSG, 10 μM), compared with non-stimulated, control plasmid (Mock)-transfected HPAECs (**Figure 3C**). Consistently, Ad*PPARγ* overexpression with RSG treatment increases HPAEC *ETV2* expression in normoxic conditions and restores ETV2 levels in hypoxic conditions (**Figure 3D**). Adenovirus-mediated-*PPARγ* overexpression in HPAECs also mitigated the hypoxia-induced increase in EndoMT markers and the reduced expression of EC markers (**Figure 3E**). Importantly, lung tissue isolated from endothelial-targeted *PPARγ* overexpressing (ePPARγOX) mice (**Figure 3F**) showed enhanced expression of both *Etv2* (**Figure 3G**) and EC markers (**Figure 3H**) with reduced expression of EndoMT markers (**Figure 3H**). To further determine the relationship between PPARγ and ETV2, we performed a series of loss of function studies. First, HPAECs were treated with *PPARγ* siRNA causing ~60% knockdown of *PPARγ* (**Figure 4A**). Reduction of *PPARγ* effectively attenuated the level of *ETV2* mRNAs (**Figure 4B**), whereas *PPARγ* expression did not change in HPAECs transfected with *ETV2* siRNA (**Figure 4C**). Similarly, in endothelial-targeted *PPARγ* knockout (ePPARγKO) mice, loss of *PPARγ* in ECs (**Figure 4D**) reduced *ETV2* expression (**Figure 4E**) and increased mesenchymal and reduced endothelial markers (**Figure 4F**) in lung homogenates *in vivo.* Collectively, these findings indicate that PPARγ functions as a direct upstream regulator of ETV2 expression in ECs.

**Figure 3.**
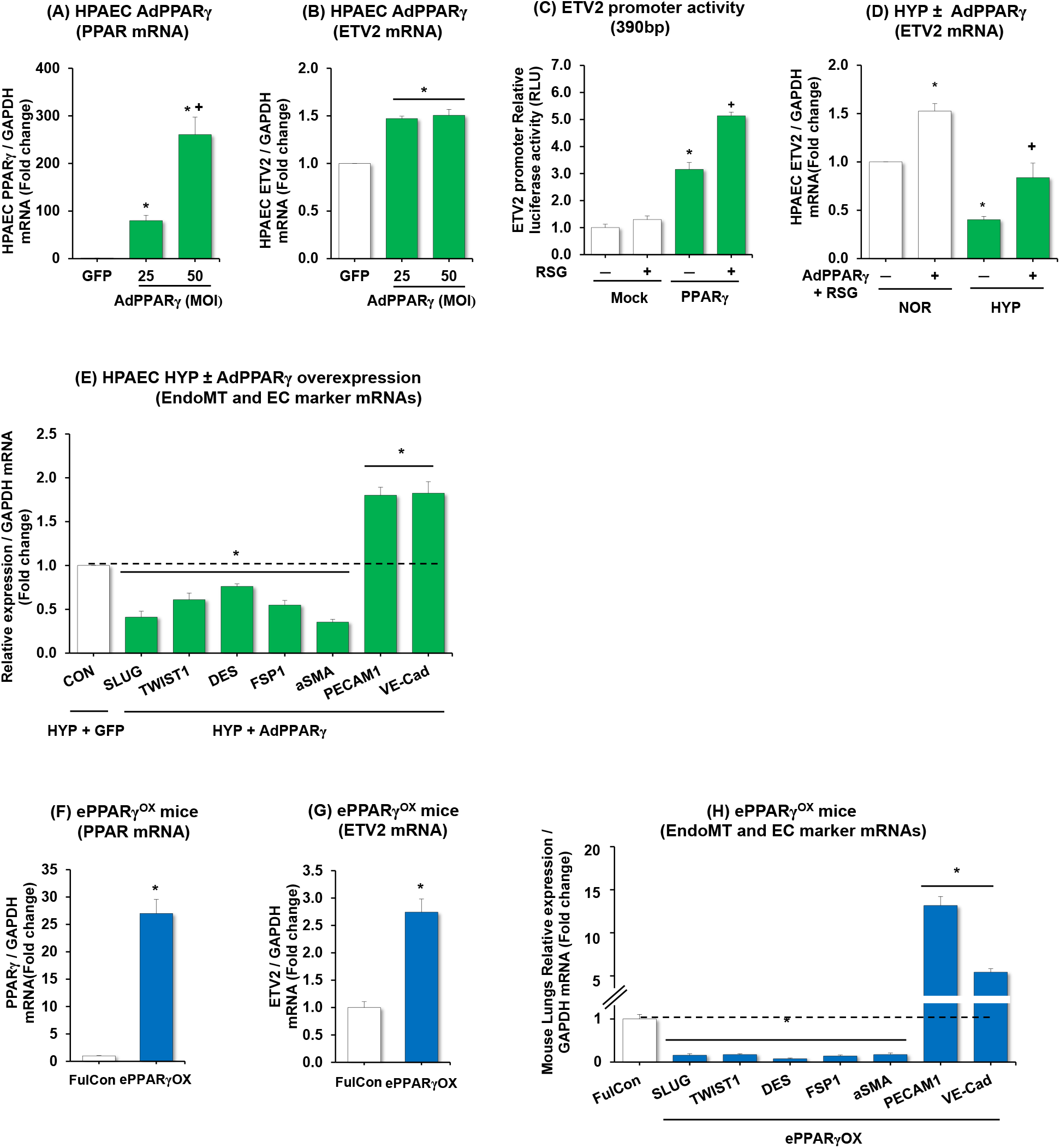
PPARγ activates ETV2 expression in HPAECs and PPARγ gain-of-function increases ETV2, PECAM1, and VE-Cad expression and attenuates levels of EndoMT markers. **(A, B)** HPAECs treated with AdPPARγ (0-50 MOI) or green fluorescent protein (GFP) constructs were subjected to qRT-PCR analysis. **(C)** PPARγ expressing plasmid (PPARγ) or control plasmid (Mock) was co-transfected with ETV2 promoter-Luc plasmid and treated with dimethyl sulfoxide (RSG/-) or rosiglitazone (RSG/+) (10 μM), then incubated for 72 hours. Fire fly luciferase activity was normalized by *Renilla* luciferase activity (right panel). **(D)** HPAECs were treated with AdPPARγ+ RSG for 6 hours, then incubated with fresh medium for an additional 72 hours under normoxic (NOR) or hypoxic (HYP) condition. RGS was treated for last 24 hours. (**E**) HPAECs treated with AdPPARγ or green fluorescent protein (GFP) constructs were cultured under hypoxic condition and subjected to qRT-PCR analysis. Each bar represents mean ± SE PPARγ, ETV2, EndoMT or EC markers level relative to GAPDH expressed as fold-change vs cells treated with GFP. n=5-6, *p<0.05 vs GFP or HYP/AdPPARγ(-). +p<0.05 vs HYP/ AdPPARγ(-). **(F-H)** whole lungs were collected from littermate control (FulCon) or endothelial-targeted PPARγ overexpression (ePPARγ^OX^) mice. Levels of lung *PPARγ* (F) or *ETV2* (G) or EndoMT and EC markers (H) were measured with qRT-PCR and expressed relative to *GAPDH* mRNA *p<0.05 vs FulCon, n=6.

**Figure 4.**
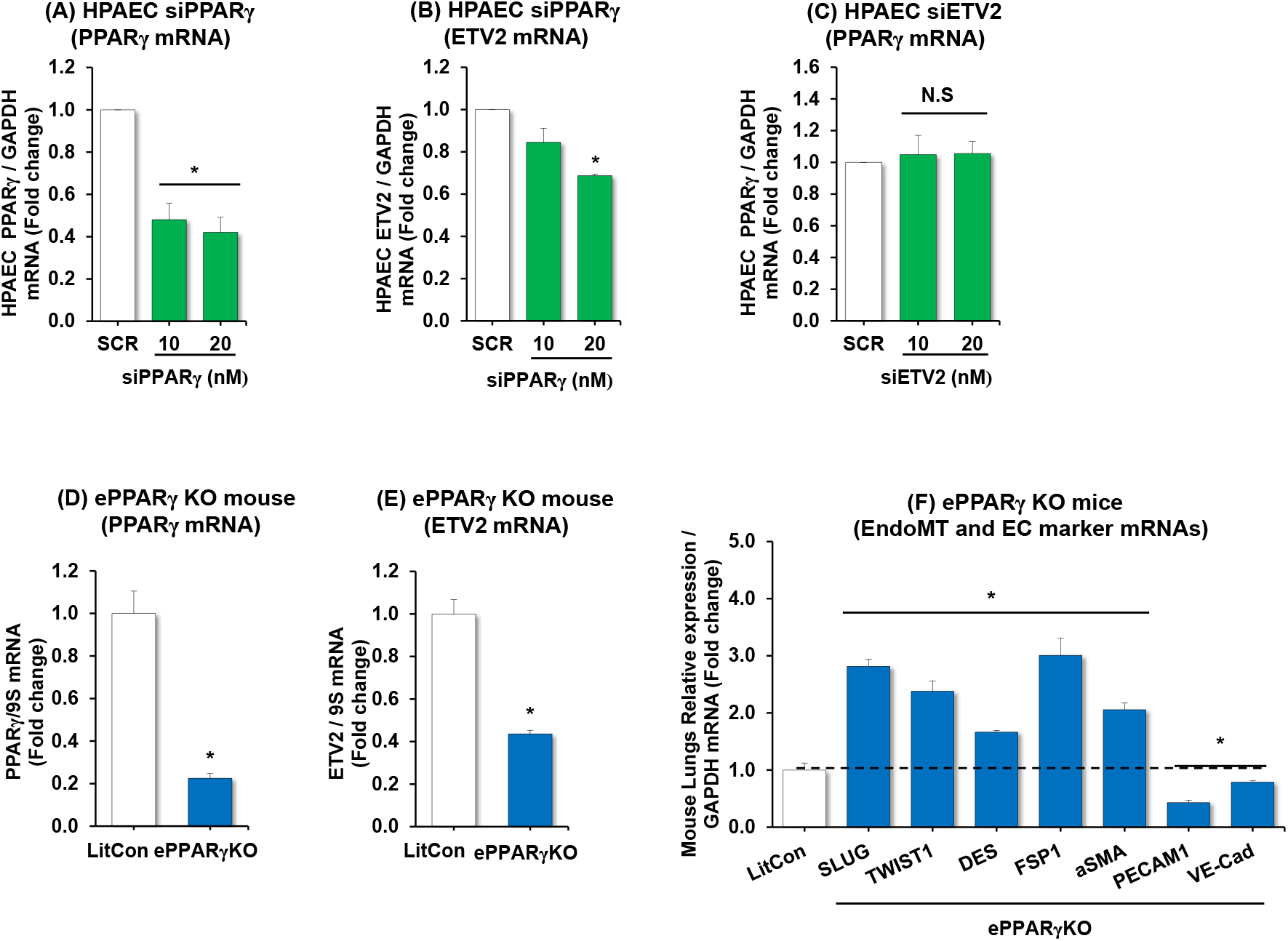
Endothelial depletion of PPARγ reduces ETV2, PECAM1, and VE-Cad expression and increases levels of EndoMT markers in mouse lungs. **(A, B)** HPAECs were treated with scrambled (SCR) or PPARγ (10-20 nM) siRNAs **(A, B)** or ETV2 (10-20 nM) siRNAs **(C)** for 6 hours, washed with fresh medium, then incubated for an additional 72 hours. Each bar represents mean ± SE *PPARγ* or *ETV2* level relative to *GAPDH* expressed as fold-change vs cells treated with SCR. n=3, *p<0.05 vs SCR. **(D-F)** Whole lungs were collected from littermate control (LitCon) or endothelial-targeted PPARγ knockout *(ePPARγ^KO^)* mice. Lung levels of *PPARγ* (D), *ETV2* (E), or EndoMT and EC markers (F) mRNA levels were measured using qRT-PCR in littermate control (LitCon) or endothelial-targeted PPARγ knockout *(ePPARγ^KO^)* mice and expressed relative to *GAPDH* mRNA *p<0.05 vs LitCon, n=6.

### *In vitro* model of EndoMT

To further investigate the role of ETV2 in EndoMT, we refined a model to induce EndoMT (termed induced-EndoMT, ***i*-EndoMT**) in HPAEC *in vitro* (30). HPAECs were treated with inflammatory mediators (IL-1β, TNFα, and TGFβ) in time-/dose-ranging regimens (Figure E2 in the online data supplement). **Figure 5A** illustrates that these inflammatory mediators, which are increased in PH, caused HPAEC to lose cobblestone morphology and become elongated and spindle-shaped, consistent with EndoMT. Under these conditions, the expression of *PPARγ*, *ETV2*, and EC markers was decreased, but EndoMT markers were increased (**Figure 5B-J**). In collagen gel contraction assays, *i*-EndoMT cells displayed enhanced gel contraction (dashed line in photomicrograph) to a fraction of the control gel contour (line graph) consistent with mesenchymal rather than endothelial functional characteristics (**Figure 5K)**. Similarly, wound healing assays demonstrated that *i*-EndoMT promoted cell migration into a monolayer wound, which is also consistent with mesenchymal cell function (**Figure 5L**). These findings suggest that previously described upregulated pro-inflammatory pathways in PH are sufficient to reduce endothelial PPARγ and ETV2 and downstream induction of EndoMT phenotypes.

**Figure 5.**
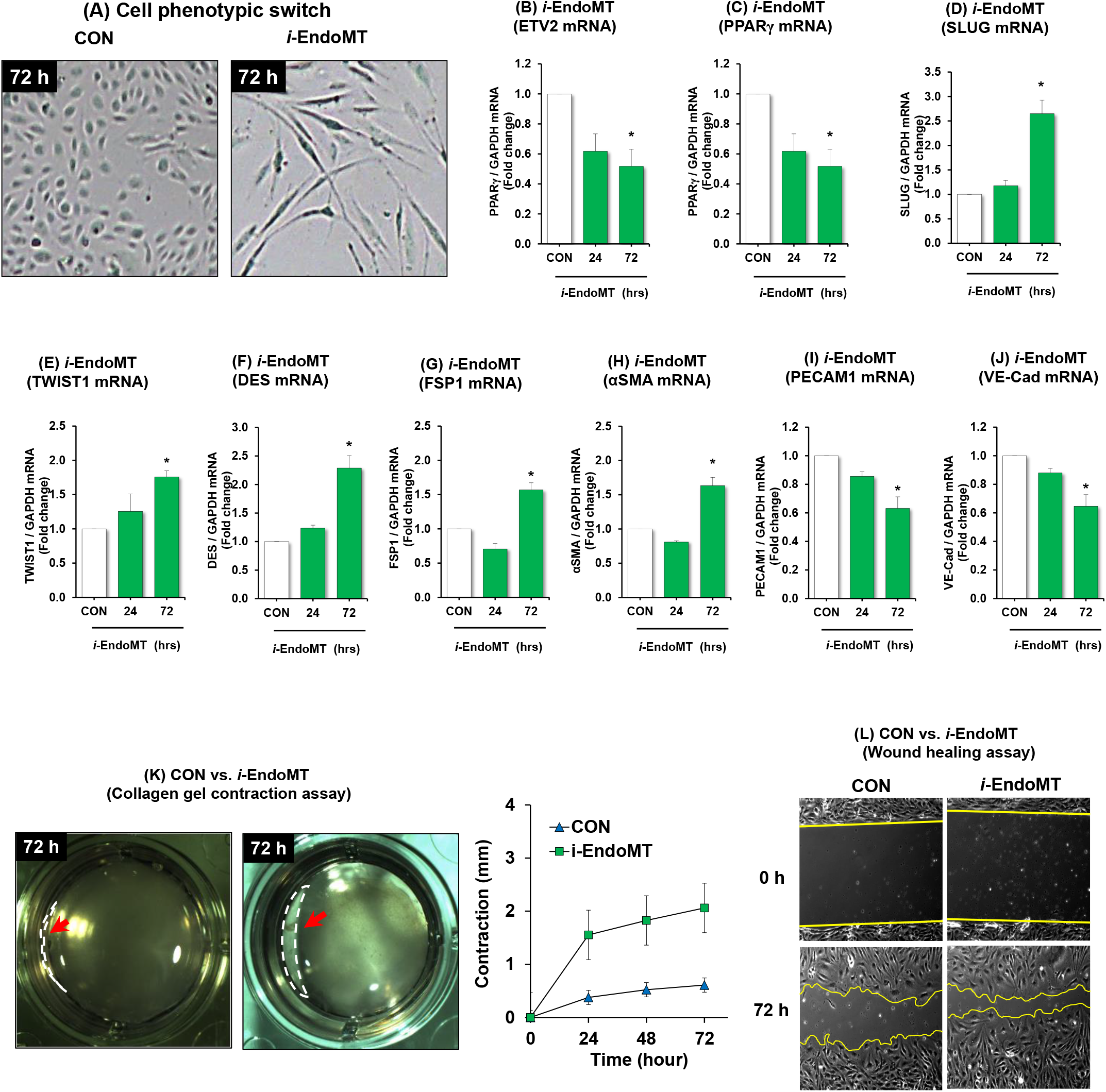
Establishment of EndoMT (*i*-EndoMT) model. **(A-J)** *i*-EndoMT cells were induced by addition of 0.1 ng/mL interleukin-1 beta (IL-1β), 10 ng/mL tumor necrosis factor alpha (TNFα), and 10 ng/mL transforming growth factor beta (TGFβ) to HPAECs up to 72 hours. Mean HPAEC *PPARγ, ETV2,* EndoMT or EC marker levels were measured with qRT-PCR. All bars represent the mean *PPARγ, ETV2* or *SLUG* mRNA levels relative to *GAPDH* ± SE expressed as fold-change vs. NOR. *p<0.05 vs. NOR, n=3. **(K)** Cell contraction assay. CON or i-EndoMT cells were harvested at 72 hours after the treatment and resuspended with a total of 5×10^5^ cells in a 1:4 ratio of cell suspension and collagen mixture provided in the cell contraction assay (Cell Biolabs). The gels were imaged at 0 hour and 72 hours post-incubation and analyzed by ImageJ software. n=2-3. Red arrow (dashed line in photomicrograph) indicates the cell contraction. **(L)** Wound healing assay. CON or i-EndoMT cells harvested at 72 hours post-treatment were harvested and replated in a wound healing chamber. Images were taken 5 hours later.

### ETV2 overexpression partially reverses *i-*EndoMT and IPAH lung fibroblasts

To further explore the ability of ETV2 to reverse EndoMT, PAECs were infected with lentiviral particles of *ETV2* undergoing *i-*EndoMT. As shown in **Figure 6A**, inducible overexpression of *ETV2* leads to a significantly reduced number of cells with mesenchymal morphology, compared to control. Cells overexpressing *ETV2* under *i-*EndoMT had decreased expression of mesenchymal markers with increased EC makers, compared to control (**Figure 6B and C**). Similar results were found when *PPARγ* was overexpressed (**Figure 6D**). More importantly, overexpression of *ETV2* converted lung fibroblasts derived from IPAH patients to cells displaying EC features (**Figure 6E-G**). Hypoxia/Sugen exposure is associated with impaired endothelium-dependent relaxation of pulmonary arteries (31). Based on hypoxia-induced decreases in ETV2 and increases in EndoMT markers, we examined whether restored ETV2 expression can improve endothelium-dependent relaxation of pulmonary arteries isolated from rats treated with the hypoxia/Sugen protocol. Overexpression of ETV2 using an adenovirus showed a strong tendency of enhanced vascular relaxation, compared to controls (**Figure 6I**). These results suggest that the *ETV2* is previously undescribed direct downstream target of PPARγ plays important functions in EndoMT and thus potentially PH pathogenesis.

**Figure 6.**
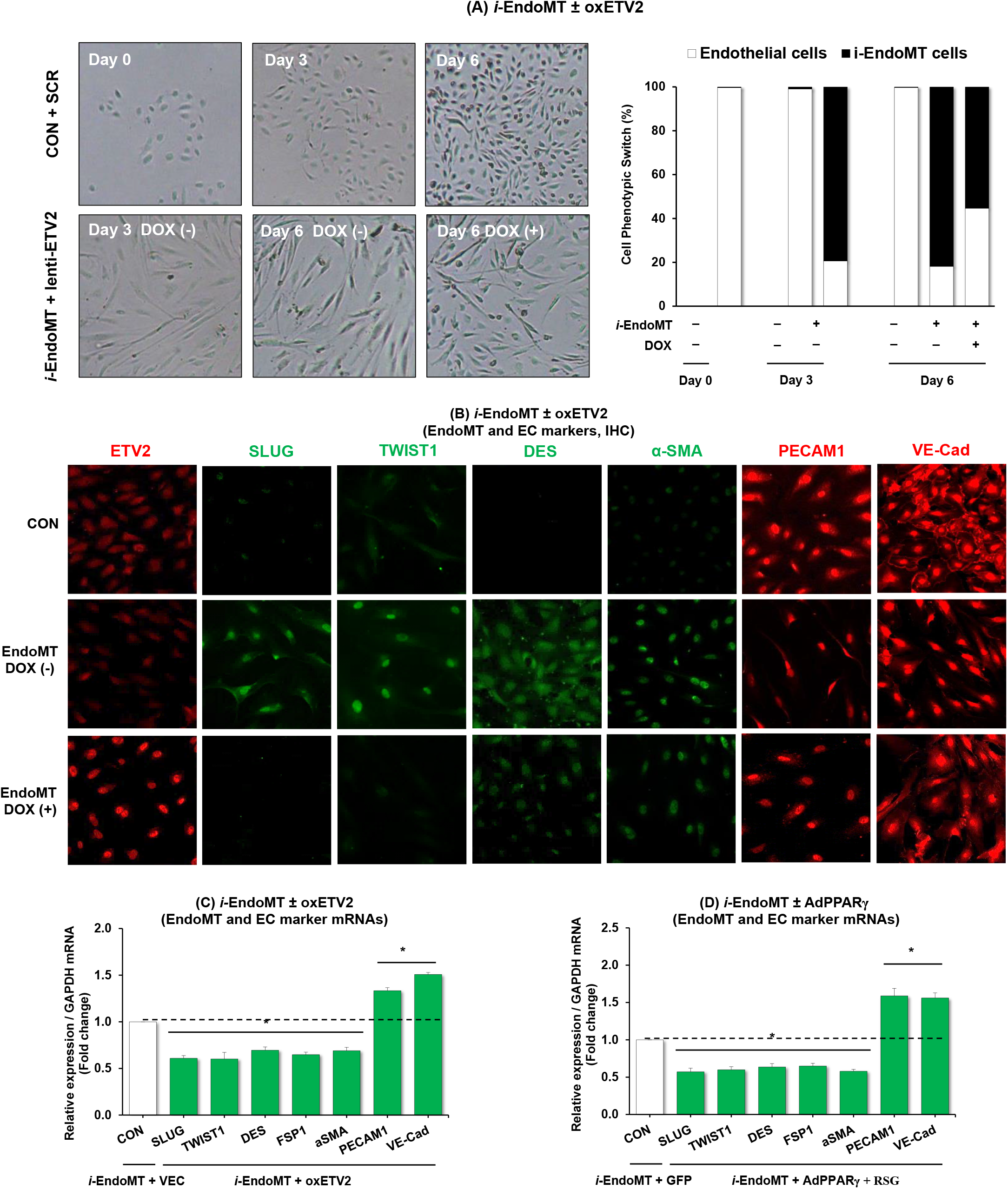
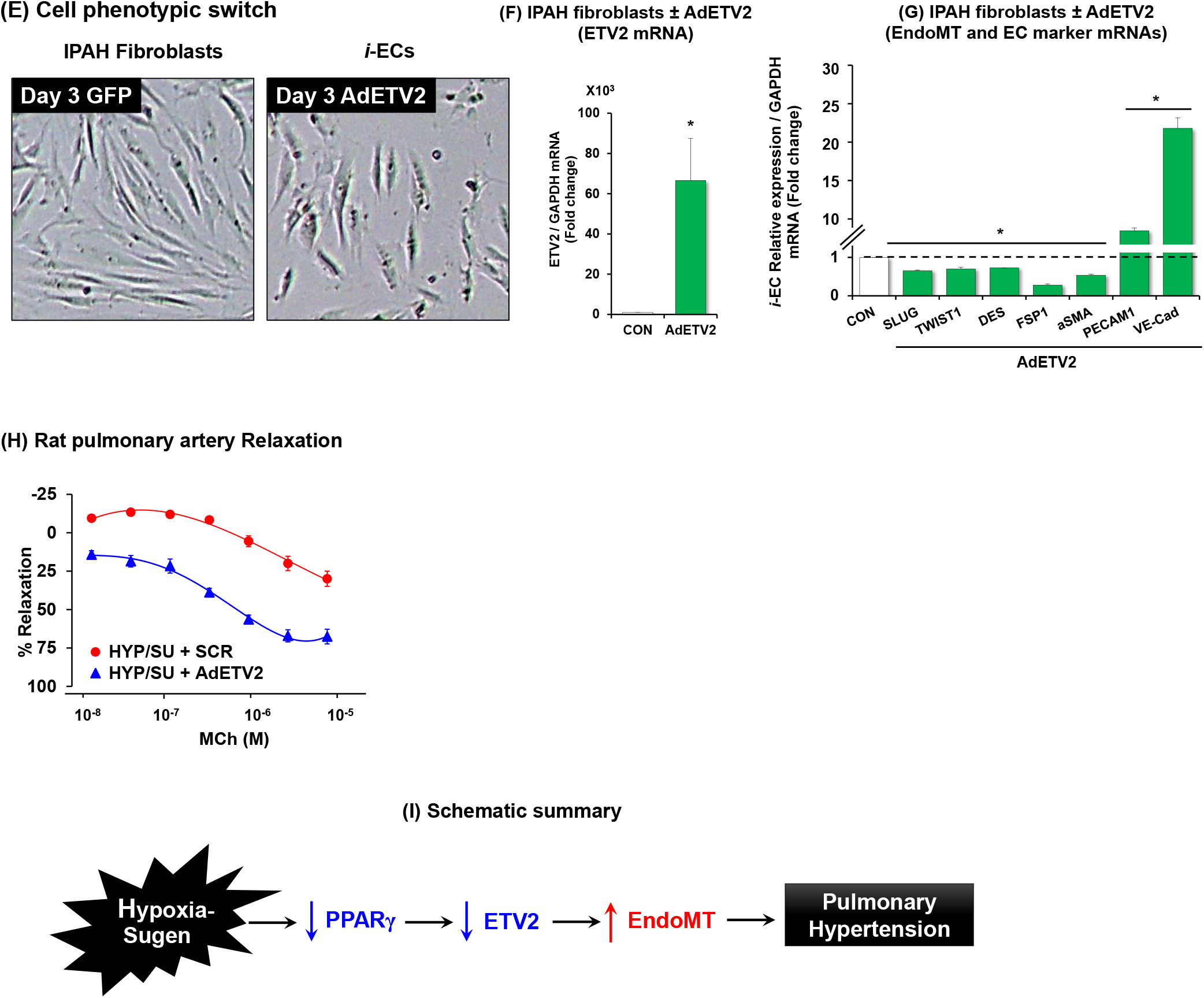
Overexpression of ETV2 induces endothelial cell phenotype and promotes vascular relaxation. **(A-B)** HPAECs infected lentiviral particles of doxycycline-inducible *ETV2* (lenti-ETV2) were incubated with 0.1 ng/mL IL-1β, 10 ng/mL TNFα, and 10 ng/mL TGFβ for 72 hours. The resulting cells were then treated ± Dox up to 6 days. A representative image of the resulting cells was shown in A (left panels). The cells were randomly selected and counted, the ratio of cobblestone morphology to elongated-spindle-shaped phenotype (A, right panel) was determined, and immunocytochemistry was performed (B). **(C, D)** HPAECs transfected with ETV2 plasmid constructs (oxETV2, 1 μg) or vector (VEC) constructs (C) or infected with AdPPARγ (25 MOI) or green fluorescent protein (GFP) constructs were incubated with 0.1 ng/mL IL-1β, 10 ng/mL TNFα, and 10 ng/mL TGFβ to HPAECs for 72 hours. RNAs from the resulting cells were subjected to qRT-PCR analysis. Twenty-four hours before cell harvest (D), the cells were treated with rosiglitazone (RSG, 10 μM). Each bar represents the mean ± SE EndoMT or EC marker level relative to *GAPDH* as indicated. *p<0.05 vs. CON/VEC or *i*-EndoMT/GFP. n=3-4. **(E-G)** Lung fibroblasts of IPAH patients were infected with AdETV2 and 3 days later, the resulting cells were imaged (E) and subjected to qRT-PCR analysis (F, G). The results are presented as fold-change vs. CON. *p<0.05 vs NOR, n=5. **(H)** Secondary branch pulmonary artery segments (3 mm in length), were isolated from hypoxia/sugen-treated Sprague-Dawley rat lungs (age 8-10 weeks) for 3 weeks and reoxygenation for 2 weeks and exposed to control conditions or 1 × 10^6^ *adETV2* for 24 hours. Studies examining endothelium-dependent relaxation were carried out by measuring responses to methacholine (MCh; 1 nM to 10 μM) and sodium nitroprusside (SNP; 0.1 nM to 1 μM). Results are expressed as mean + SEM. Concentration-response curves were generated to the contractile agonist phenylephrine (PE, 0.1 nM to 10 μM) and ED80 determined. n=5-8. **(I)** A hypothetical schema defining the role of PPARγ/ETV2 on EndoMT in PH pathogenesis.

## DISCUSSION

The current study provides several novel observations that could advance understanding of PH pathogenesis; 1) ETV2 is downregulated in lungs and PAECs from patients with IPAH, in lungs of mice with PH caused by HYP/Su, in hypoxia-exposed HPAECs, and in *i*-EndoMT cells. 2) Reduction of ETV2 (i.e., *Etv2^+/-^* mice) leads to a spontaneous development of PH, augmented RVH, and increased expression of EndoMT markers. 3) PPARγ acts as an upstream regulator of ETV2 expression. 4) Sustained expression of PPARγ or ETV2 in ECs inhibits the progression of EndoMT. 5) ETV2 is able to revert IPAH lung myofibroblasts to a more EC-like phenotype. Collectively, these findings suggest that the PPARγ-ETV2 axis and the regulation of EndoMT play an important role in PH **(Figure 6I**).

PH is a chronic cardiopulmonary disorder that causes significant morbidity and mortality (1, 3, 32, 33). Current targeted therapies in PH fail to reverse pulmonary vascular remodeling and are frequently not recommended for patients with more common forms of PH (34, 35). These observations indicate that a greater understanding of PH pathogenesis may permit more effective therapeutic strategies. Recent studies using cell lineage tracking analysis have shown that EndoMT contributes to the initiation and progression of PH (5, 9, 30). This role of EndoMT was further supported by the colocalization of von Willebrand factor and αSMA in pulmonary endothelium from HYP/Su rodent models and PAH patients (30). However, the molecular mechanisms underlying EndoMT in the pathogenesis of PH remain largely unknown. In this study, we propose the PPARγ-ETV2 axis as a key mechanism in regulating EndoMT in PH.

ETV2 functions as a master regulator for EC generation and function (10, 36). While deficiency in *Etv2* leads to a complete lack of vascular ECs (11, 12), forced expression of ETV2 is sufficient to reprogram non-ECs including mesenchymal cells such as human dermal fibroblasts into functional ECs (17, 18). These findings suggested that substantial loss of EC in the progression of PH could be due to reduced expression of ETV2. In agreement with this hypothesis, we found a significant reduction in ETV2 expression in clinical and experimental PH lung samples. Reduction of ETV2 (*Etv2^+/-^* mice) was sufficient to induce spontaneous PH and RVH in mice as well as upregulated expression of EndoMT makers and concomitant reductions of EC markers. These results strongly suggest the functional significance of ETV2 in PH.

Although the precise molecular mechanisms by which ETV2 regulates PH is unknown, it is clear that ETV2 acts as a direct transcriptional regulator of diverse EC genes including CDH5 and CD31 (37). In addition, a recent report showed that ETV2 can regulate the expression of Robo4, one of the critical EC genes via DNA methylation (38). Further, we showed that ETV2 functions with valproic acid, a histone acetylation modifier in direct cell reprogramming (18). These findings suggest that decreased expression of ETV2 in PH induces inactivation of EC gene expression and loss of normal EC function. Currently, it is not known whether the increase of mesenchymal cells in PH is a direct outcome of ETV2 downregulation or secondary to the loss of ECs. The molecular mechanisms of ETV2-EndoMT remain an area of active investigation in our laboratories.

The current study extends the field by providing novel insights into the regulation of ETV2 expression in PH. Given the established role of PPARγ in PH pathobiology, we performed *in silico* analysis of the upstream promoter region of ETV2 and identified several putative PPARγ binding sites. Our results showed that PPARγ can transcriptionally activate ETV2 expression. This finding was further supported by a series of experiments *in vivo* and *in vitro.* Mice lacking endothelial *Pparγ* exhibited a reduced level of expression of *Etv2* in lungs, while mice overexpressing endothelial *Pparγ* showed augmented level of *Etv2.* Overexpression or knockdown experiments *in vitro* further substantiated these results. Collectively, these data suggest that PPARγ regulates PH in part through direct transcriptional activation of ETV2. Therefore, further studies to determine the functional consequences of the PPARγ-ETV2 axis in PH in vivo will be warranted.

One of the key findings of this study was to refine *in vitro* model which can mimic EndoMT *in vivo.* A recent study demonstrated that a combination of inflammatory mediators (IL-1β, TNFα, and TGFβ) induced EndoMT (30). The current study extends this report to demonstrate that these three inflammatory mediators phenotypically switched not only endothelial cells (cobblestone morphology) to mesenchymal cells (elongated, spindle-shaped myofibroblastic cell morphology) *in vitro*, but also induced functional alterations including enhanced contractile and migratory phenotypes in *i*-EndoMT cells. Consistent with these phenotypic and functional features, *i*-EndoMT cells expressed higher levels of EndoMT markers with downregulated levels of *ETV2* and *PPARγ.* Importantly, we found that overexpression of *PPARγ* or *ETV2* was able to reduce the expression of mesenchymal markers in cells undergoing *i*-EndoMT, suggesting the potential role of PPARγ and ETV2 in EndoMT.

Recent studies have shown that ETV2 can directly reprogram non-ECs into ECs (17, 18). In addition, delivery of lentiviral *ETV2* into ischemic hindlimbs or infarcted heart promotes recovery of ischemic damage as evidenced by enhanced perfusion and cardiac functions, respectively (16, 39). Interestingly, massive neovascularization also was accompanied in the damaged tissues upon the overexpression of ETV2, suggesting its potent function in endothelial reprogramming and functional recovery of vascular defects. In this study, we showed that PAH-derived lung myofibroblasts acquire EC characteristics while losing a mesenchymal phenotype when *ETV2* was overexpressed. Further, overexpression of ETV2 showed improved vascular relaxation in rat pulmonary arteries treated with the hypoxia/Sugen. These results suggest that ETV2 could be a therapeutic target for PH in potentially reverting mesenchymal cells to functional ECs.

In summary, to our knowledge, the current study provides for the first time uncovering novel functions of the PPARγ/ETV2 axis in regulating EndoMT during the pathogenesis of PH. Thus, the outcome of the proposed studies will advance current knowledge on EndoMT in PH and provide a new therapeutic paradigm for treating PH patients.

## Acknowledgments

This study was supported by funding from VA BLR&D Merit Review Award (I01 BX004263 to CMH), NIH National Heart, Lung, and Blood Institute R01 grants (HL102167 to CMH and RLS, HL119291 to CP, and HL133053 to BYK). Data/Tissue samples provided by PHBI under the Pulmonary Hypertension Breakthrough Initiative (PHBI). Funding for the PHBI is provided under an NHLBI R24 grant, #R24HL123767, and by the Cardiovascular Medical Research and Education Fund (CMREF).

## Author disclosures

The authors have no competing interests. The contents of this report represent the views of the authors and do not represent the views of the Department of Veterans Affairs or the United States Government.

## Notes

### Competing Interest Statement

The authors have declared no competing interest.

