## Supplementary Methods-Figure Legends-Figures for "PPARγ/ETV2 Axis Regulates Endothelial-to-Mesenchymal Transition in Pulmonary Hypertension"

### Online Supplemental Data

### PPARγ/ETV2 Axis Regulates Endothelial-to-Mesenchymal Transition in Pulmonary Hypertension

Dong Hun Lee[a,d,†] , Minseong Kim[a,l,†], Sarah S. Chang[b,c, †], Andrew J. Jang[e], Juyoung Kim[a,d], Jing Ma[b,c], Michael J. Passineau[e], Raymond L. Benza[f], Harry Karmouty-Quintana[g,h], Wilbur A. Lam [i], Roy L. Sutliff[b,c,j], C. Michael Hart[b,c] , Changwon Park[a,k,†,*], and Bum-Yong Kang[a,b,c,*]

[a]Department of Pediatrics, Division of Hematology, Oncology, and BMT, [b]Department of Medicine, Division of Pulmonary, Allergy, Critical Care, and Sleep Medicine, Emory University School of Medicine, and [c]Atlanta Veterans Healthcare System, Decatur, GA,

[d]Department of Biological Sciences, Chonnam National University, 77 Yongbong-ro, Buk-gu, Gwangju, 61186, Republic of Korea

[e]Cardiovascular Institute, Department of Medicine, Allegheny Health Network, Pittsburgh, PA.

[f]The Ohio State University Wexner Medical Center, Columbus, Ohio,

[g]Department of Biochemistry and Molecular Biology, [h]Divisions of Critical Care & Pulmonary and Sleep Medicine, Department of Internal Medicine, McGovern Medical School, University of Texas Health Science Center at Houston, Houston, TX

[i]Georgia Institute of Technology, Atlanta, GA, USA.

[j]National Heart, Lung and Blood Institute. National Heart, Lung, and Blood Institute. Bethesda, MD, USA.

### SUPPLEMENTARY FIGURE LEGENDS

**Supplementary Figure E1. PPARγ is reduced in lungs and PAECs isolated from IPAH patients, lungs of hypoxia/sugen-treated mice, and in hypoxia-exposed HPAECs**

**(A-B)** The expression of lung or PAEC PPARγ was measured with qRT-PCR and expressed relative to *GPADH* mRNA ± SE as fold-change vs. CON. *p<0.05 vs NOR, n=5. **(C)** Whole lung or pulmonary artery homogenates were collected from mice under normoxic (NOR, 21%)/sugen or hypoxic (HYP, 10% $O_2$)/sugen mice for 3-weeks. Lung PPARγ expression was measured with qRT-PCR and expressed relative to lung *Gapdh* mRNA *p<0.05 vs NOR, n=6. **(D)** HPAECs were exposed with normoxia (NOR, 21% $O_2$) or hypoxia (HYP, 1% $O_2$) for 72 hours. Mean HPAEC PPARγ expression was measured with qRT-PCR. All bars represent the mean *PPARγ* mRNA levels relative to *GAPDH* ± SE expressed as fold-change vs. NOR. *p<0.05 vs. NOR, n=6.

**Supplementary Figure E2. Inflammatory mediators reduce PPARγ and ETV2 levels and induce EndoMT maker SLUG in HPAECs**

HPAECs were dose-dependently treated with interleukin-1 beta (IL-1β), tumor necrosis factor alpha (TNFα), or transforming growth factor beta (TGFβ) for 72 hours. **(Upper left panels**) A representative image of *i*-EndoMT cells with the individual factors (0.1 ng/mL IL-1β, 10 ng/mL TNFα**,** or 10 ng/mL TGFβ). **(Upper right and lower panels)** HPAECs cultured with different concentrations of the factors were subjected to qRT-PCR analysis. Mean HPAEC *PPARγ*, *ETV2*, and *SLUG* expression were measured with qRT-PCR. All bars represent the mean *PPARγ, ETV2* or *SLUG* mRNA levels relative to *GAPDH* ± SE expressed as fold-change vs. NOR. *p<0.05 vs. NOR, n=3.

### Supplementary Figures

### PPARγ/ETV2 axis Regulates Endothelial-Mesenchymal Transition in Pulmonary Hypertension

Dong Hun Lee[a,d,†] , Minseong Kim[a,l,†], Sarah S. Chang[b,c, †], Andrew J. Jang[e], Juyoung Kim[a,d], Jing Ma[b,c], Michael J. Passineau[e], Raymond L. Benza[f], Harry Karmouty-Quintana[g,h], Wilbur A. Lam [i], Roy L. Sutliff[b,c,j], C. Michael Hart[b,c] , Changwon Park[a,k,†,*], and Bum-Yong Kang[a,b,c,*]

[a]Department of Pediatrics, Division of Hematology, Oncology, and BMT, [b]Department of Medicine, Division of Pulmonary, Allergy, Critical Care, and Sleep Medicine, Emory University School of Medicine, and [c]Atlanta Veterans Healthcare System, Decatur, GA,
[d]Department of Biological Sciences, Chonnam National University, 77 Yongbong-ro, Buk-gu, Gwangju, 61186, Republic of Korea
[e]Cardiovascular Institute, Department of Medicine, Allegheny Health Network, Pittsburgh, PA.
[f]The Ohio State University Wexner Medical Center, Columbus, Ohio,
[g]Department of Biochemistry and Molecular Biology, [h]Divisions of Critical Care & Pulmonary and Sleep Medicine, Department of Internal Medicine, McGovern Medical School, University of Texas Health Science Center at Houston, Houston, TX
[i]Georgia Institute of Technology, Atlanta, GA, USA.
[j]National Heart, Lung and Blood Institute. National Heart, Lung, and Blood Institute. Bethesda, MD, USA.
[k]Department of Cellular and Molecular Physiology, Louisiana State University Health Science Center, Shreveport, LA

### Supplementary Fig. E1

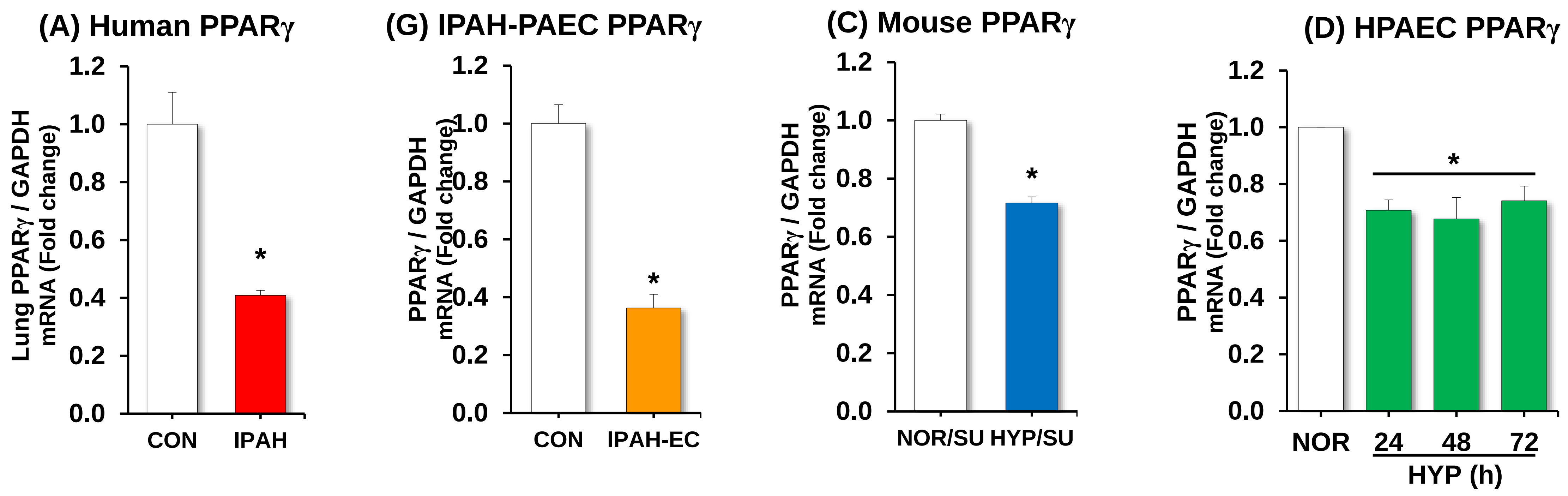

### Supplementary Fig. E2

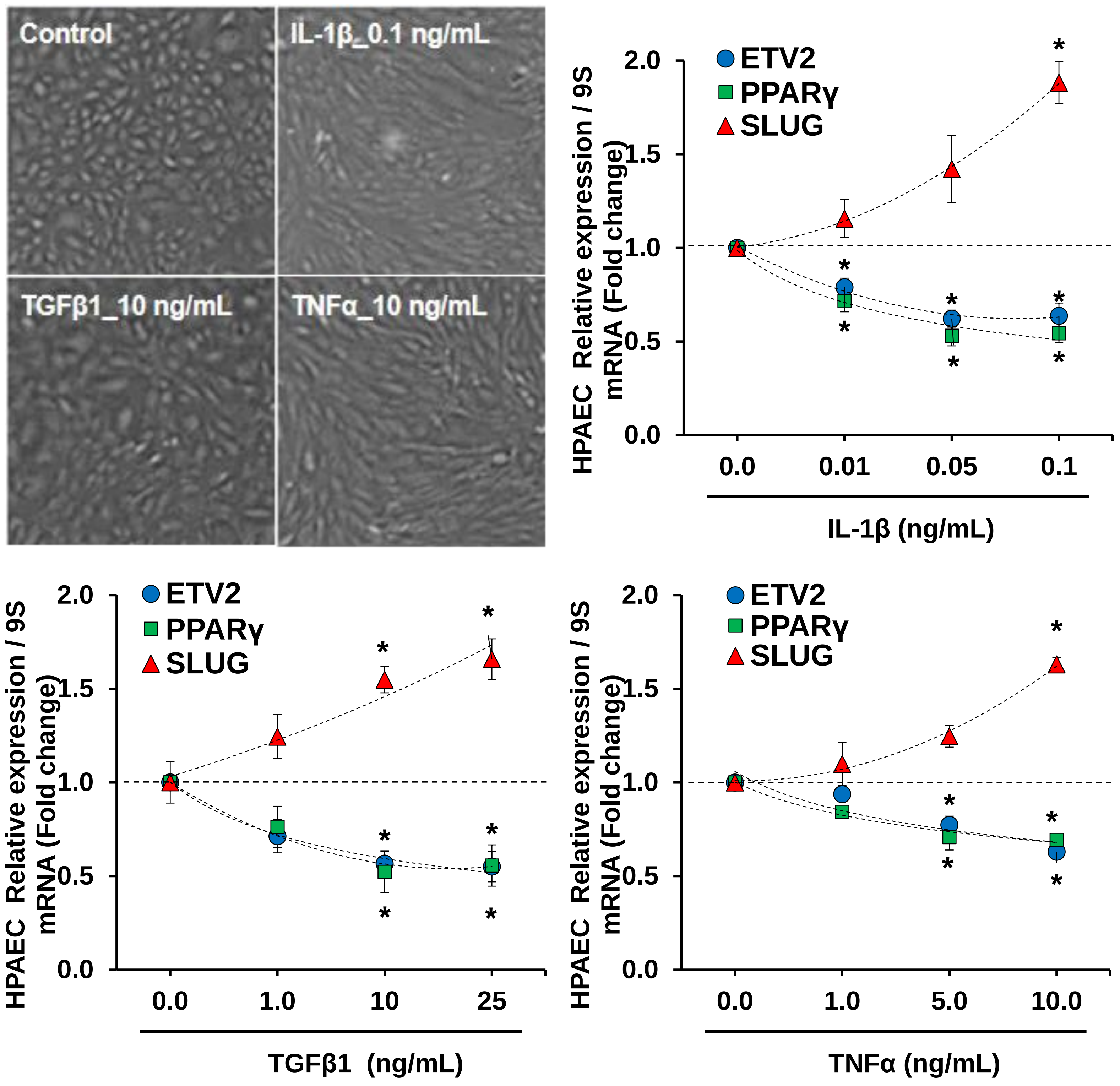
